# Magnetic Resonance Fingerprinting based Thermometry (MRFT): application to ex vivo imaging near DBS leads

**DOI:** 10.1101/2023.01.11.523421

**Authors:** Enlin Qian, Pavan Poojar, Maggie Fung, Zhezhen Jin, Thomas Vaughan, Devashish Shrivastava, David Gultekin, Tiago Fernandes, Sairam Geethanath

**Affiliations:** Columbia Magnetic Resonance Research Center, Columbia University, New York, NY, United States; Department of Biomedical Engineering, Columbia University, New York NY, United States; Department of Biostatistics, Columbia University, New York, NY, United States; Accessible MR Laboratory, Biomedical Engineering and Imaging Institute, Dept. of Diagnostic, Molecular and Interventional Radiology, Icahn School of Medicine at Mt. Sinai, New York, NY, United States; GE Healthcare, New York, NY, 10065, USA; ISR - Lisboa/LARSyS and Department of Bioengineering, Instituto Superior Técnico – Universidade de Lisboa, Lisbon, Portugal

## Abstract

The purpose of this study is to demonstrate the first work of T_1_-based magnetic resonance thermometry using magnetic resonance fingerprinting (dubbed MRFT). We compared temperature estimation of MRFT with proton resonance frequency shift (PRFS) thermometry on *ex vivo* bovine muscle. We demonstrated MRFT’s feasibility in predicting temperature on *ex vivo* bovine muscles with deep brain stimulation (DBS) lead. B_0_ maps generated from MRFT were compared with gold standard B_0_ maps near the DBS lead.

All experiments were performed on a 3 Tesla whole-body GE Premier system equipped with a 21-channel receive head coil (GE Healthcare, Milwaukee, WI). Four fluoroptic probes were used to measure the temperature at the center of a cold muscle (probe 1), the room temperature water bottle (probe 2), and the center and periphery of the heated muscle (probes 3 and 4). We selected regions of interest (ROIs) around the location of the probes and used simple linear regression to generate the temperature sensitivity calibration equations that convert T_1_ maps and Δs maps to temperature maps. We then repeated the same setup and compared MRFT, PRFS thermometry temperature estimation with gold standard probe measurements. For the MRFT experiment on DBS lead, we taped the probe to the tip of the DBS lead and used a turbo spin echo (TSE) sequence to induce heating near the lead. We selected ROIs around the tip of the lead to compare MRFT temperature estimation with probe measurements. Vendor-supplied B_0_ mapping sequence was acquired to compare with MRFT-generated B_0_ maps.

We found strong linear relationships (R^2^>0.958) between T_1_ and temperature and Δs and temperatures in our temperature sensitivity calibration experiment. MRFT and PRFS thermometry both accurately predict temperature (RMSE<1.55 °C) compared to probe measurements. MRFT estimated temperature near DBS lead has a similar trend as the probe temperature. Both B_0_ maps show inhomogeneities around the lead.

## Introduction

The clinical development of deep brain stimulation (DBS) is considered one of the most significant advances in the field of neuroscience in the past two decades (Lozano *et al* 2019). DBS is a neurosurgical procedure that involves implanting electrodes into certain areas of the brain and generating electric impulses that directly intervene in pathological neural circuits. After the implantation of the DBS system, patients may require MRI to monitor and evaluate a wide range of disease processes (Martin 2019). MRI is used for presurgical evaluation including trajectory planning as well as the selection of optimal location and postsurgical procedures such as postoperative target confirmation for detecting any procedural hemorrhage postoperatively and for optimizing subsequent implantable pulse generator (IPG) programming (Gupte *et al* 2011). During MRI scans, the time-varying electromagnetic (EM) field induces a current in DBS lead. The current dissipates in the surrounding tissue and produces localized heating (Shrivastava *et al* 2012). Such temperature change can result in thermal damage and tissue necrosis (Gupte *et al* 2011). However, non-invasive temperature assessment and standardized temperature measurement remain challenging (Quinn 2013, Nabavi 2010).

Non-invasive imaging techniques such as magnetic resonance imaging (MRI), computed tomography (CT), ultrasound, high-intensity focused ultrasound (HIFU), photoacoustic imaging, microwave radiometry, near-infrared spectroscopy (NIRS), etc. are used for internal temperature measurement (Raiko *et al* 2020). Compared to other imaging techniques, MRI is found to be suitable for non-invasive thermometry due to the spatial resolution and anatomical coverage per unit time of MRI, its lack of ionizing radiation, and the ability to provide contrasts images as well as temperature maps (Winter *et al* 2016, Rieke 2012). Most parameters that influence MRI signals are temperature-dependent (Odéen and Parker 2019).

Several MR thermometry methods have been carefully studied through the years. Chen et al. investigated phase-drift correction proton resonance frequency shift (PRFS) in a low-field MR scanner (Chen *et al* 2018). Salomir et al. developed a novel reference-free PRFS MR thermometry based on a physically meaningful formalism (Salomir *et al* 2012) and depended on reconstructing background phase. Liu et al. used a geometric model based method to correct PRFS-based MR thermometry for fat interference (Liu *et al* 2019). Challenges in PRFS-based MR thermometry include temperature errors induced by temperature-dependent magnetic susceptibility (Sprinkhuizen *et al* 2010), motion (Rieke *et al* 2007, Grissom *et al* 2010), and PRFS underestimation in adipose tissues (Odéen and Parker n.d.). Weber et al. performed thermometry near metallic devices using multispectral imaging (Weber *et al* 2017). Baron et al. studied T_1_ and T_2_ temperature dependence of female human breast adipose tissue at 1.5T (Baron *et al* 2015). By combining variable flip angle (VFA) and fast EPI readout, short scan times of T_1_-based MR thermometry have been obtained (Diakite *et al* 2014, Hey *et al* 2012, Todd *et al* 2014). T_1_-based MR thermometry methods have challenges such as long acquisition time and the requirement of a 180^0^ pulse. The temperature dependence of T_1_ also varies among different tissue types.

Magnetic resonance fingerprinting (MRF) is an acquisition-reconstruction framework that simultaneously estimates T_1_, T_2,_ and other MR parameters using a highly undersampled pseudo-random acquisition schedule (Ma *et al* 2013). It has three advantages over the commonly applied PRFS-based MR thermometry: (1) T_1_-based MRF thermometry is reference-free; (2) MRF is more robust to both intra- and inter-scan motion; (3) MRF estimates multiple temperature-dependent tissue parameters such as T_1_, T_2_, PD (Ma *et al* 2013, Qian *et al* 2022, Balsiger *et al* 2018), ADC (Panda *et al* 2019), etc. and system parameters such as B_0_ (Ma *et al* 2013), B_1_ (Buonincontri and Sawiak 2016) simultaneously. These quantitative tissue and system parameters can be leveraged to obtain multiple temperature-dependent observations along with the opportunity to correct these estimates with the inclusion of system imperfections in B_0_, for example. This study demonstrates T_1_-based MRF thermometry (dubbed MRFT) and its application to DBS lead heating in an ex-vivo setting. To this end, we (1) perform temperature sensitivity calibration experiments for both MRFT and PRFS-based thermometry; (2) compare temperature maps generated using MRFT with PRFS thermometry and gold standard fluoroptic probe temperature; (3) compare MRFT temperatures and fluoroptic probe measurements surrounding DBS lead in bovine muscle. (4) compare MRF generated B_0_ map with gold standard B_0_ map surrounding DBS lead in bovine muscle.

## Methods

### Bovine muscle thermometry study

All experiments were performed on a 3 Tesla whole-body GE Premier system equipped with a 21-channel receive head coil (GE Healthcare, Milwaukee, WI). We used fluoroptic probes for the gold standard temperature measurement with a precision of 0.1 °C and a temporal resolution of 1 second (STB Medical, LumaSense Technologies, Santa Clara, CA).

Figure 1(a) shows the setup for the *ex vivo* sensitivity calibration experiment for T_1_-MRFT and PRFS thermometry methods. The sample consisted of three parts: cold muscle, a water bottle at room temperature, and heated muscle. The cold bovine muscle was stored in the refrigerator overnight before the experiment. The room-temperature water bottle was stored in the scanner room. The heated muscle was prepared in the microwave for 30 seconds before the experiment. Four fluoroptic probes were used to measure temperature at the center of a cold muscle (probe 1), the room temperature water bottle (probe 2), and the center and periphery of the heated muscle (probes 3 and 4).

**Figure 1:**
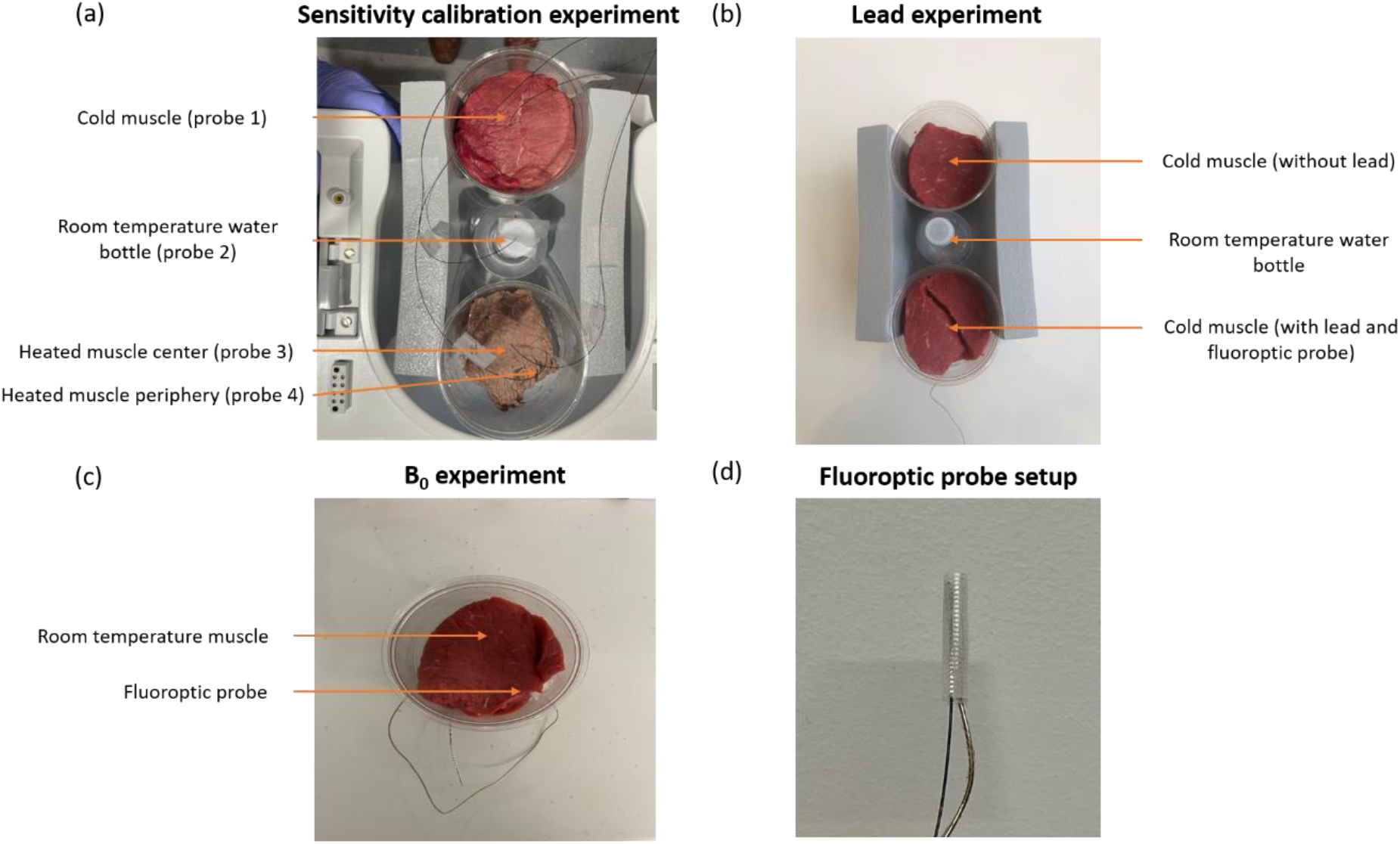
Figure 1 shows the setup for (a) calibration experiment, (b) lead experiment and (c) B0 experiment. The calibration experiment consists of three parts: cold muscle, a room temperature water bottle, and heated muscle. Four fluoroptic probes were used to measure temperature of the center of a cold muscle, the room temperature water bottle, and the center and periphery of the heated muscle. The lead experiment consists of three parts: cold muscle without lead, a room temperature water bottle, and cold muscle with lead. The lead is inserted from the bottom of the muscle and the fluoroptic probe is positioned at the tip of the lead. (c) shows the experimental setup of the B0 experiment. The DBS lead was inserted from the right bottom of the room temperature bovine muscle. (d) shows how the probe was taped to the DBS lead for the lead experiment. The fluoroptic probe was taped to the second electrode on the DBS lead.

Figure 1(b) shows the *ex vivo* DBS lead temperature experiment. The sample in this case consisted of three parts: bovine muscle at room temperature without lead, a water bottle and bovine muscle with DBS lead at room temperature. All bovine muscles were stored in the scanner room the night before the experiment. The DBS lead was inserted from the bottom of the muscle and the fluoroptic probe was positioned at the tip of the lead, as shown in Figure 1(d). We taped the fluoroptic probe to the second electrode of the lead because it has more significant temperature changes in *in vivo* experiments, as shown in previous literature (Shrivastava *et al* 2012).

Figure 1(c) shows the *ex vivo* B_0_ lead experiment. The experiment consisted of a room-temperature bovine muscle with lead inserted from the bottom. The setup was positioned in the axial orientation to accommodate the orientation limitation of the gold standard B_0_ mapping sequence.

### MRF acquisition and reconstruction

The TR/FA scheme of the MRF sequence used parameters reported in previous literature (Jiang *et al* 2015, Shridhar Konar *et al* 2021). The sequence parameters were: Field of View (FOV) of 25 cm × 25 cm, matrix size=225×225, sampling bandwidth=±250 kHz, TE=2.3 ms, slice thickness=5 mm, number of slices=1. The acquisition time was 18 seconds/slice. The raw data were reconstructed using singular value decomposition (SVD) based dictionary matching in MATLAB (MathWorks, Natick, MA). The reconstruction time was ∼30 seconds/slice.

All reconstructions were performed in line on the scanner to visualize temperature and T_1_ maps (Supplementary video 1). We compiled our scripts using Matlab Runtime 9.5 (MathWorks, Natick, MA) on a Ubuntu 20.04 system. We then transferred the compiled file to the scanner and linked the scripts to the reconstruction pipeline on the scanner.

### Temperature sensitivity calibration

#### MRFT temperature sensitivity calibration

We performed the MRFT sensitivity calibration experiment using the MRF sequence detailed in the MRF acquisition and reconstruction section. The MR protocol for the temperature calibration experiment consisted of a T_1_w turbo spin echo (TSE) structural scan, followed by an MRF sequence every five minutes for two hours. The linear relationship between temperature (T) and T_1_ can be expressed as in (Odéen and Parker 2019, Parker *et al* 1983, Parker 1984, Lewa and Majewska 1980, Bottomley *et al* 1984)

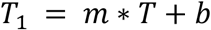

Where m and b are tissue-specific and have to be determined using calibration methods. In this study, we only considered one tissue type. We selected regions of interest (ROIs) of 25 pixels on T_1_ maps around the location of the fluoroptic probe based on the TSE images. We calculated the mean of the ROIs, and simple linear regression and R_2_ statistics were calculated using least squares fitting in the R programming language to generate the temperature sensitivity calibration equation (detailed in the Statistics section below). The x-axis for the regression curve is T_1_ values, and the y-axis is temperatures. The calibration equation converted T_1_ maps to temperature maps and was later used for generating temperature maps in the MRFT/PRFS comparison experiment and DBS lead experiment.

#### PRFS thermometry temperature sensitivity calibration

We performed the PRFS sensitivity calibration experiment using a SPoiled Gradient-Recalled (SPGR) sequence with the following parameters: TR=100 ms, TE=4.2 ms, flip angle=36°, FOV of 25 cm × 25 cm, slice thickness=5 mm, matrix size=225×225, excitation bandwidth=1.3 kHz, readout bandwidth=0.5 kHz/pixel, acquisition time=16 sec. The MR protocol for the calibration experiment consisted of a TSE structural scan, followed by an SPGR sequence every five minutes for two hours. The linear relationship between change in temperature (ΔT) and change in phase (Δϕ) can be expressed as in (Winter *et al* 2016):

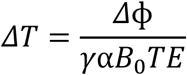

Where *γ* is the gyromagnetic ratio, α is the thermal coefficient, B_0_ is the static magnetic field and TE is the echo time of the sequence. The equation can be further simplified as:

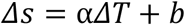

where α is the thermal coefficient and b is a constant. To effectively remove the phase wrapping during heating, the phase difference is constructed as (Peters 2000):

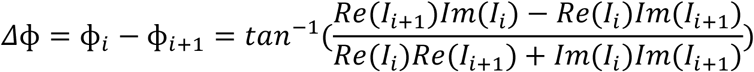

Where Re and Im are the real and imaginary parts of images I_1_ and I_2_.

We selected ROIs of 25 pixels on Δs maps around the location of the fluoroptic probe based on the TSE images. We calculated the mean of the ROIs and simple linear regression and R^2^ statistics were calculated using least squares fitting in the R programming language to generate the temperature sensitivity calibration equation (detailed in the Statistics section below). The x-axis for the regression curve is Δs, and the y-axis is temperatures. The calibration equation converted Δs to temperature maps and was later used for generating temperature maps in the MRFT/PRFS comparison experiment. To validate our PRFS thermometry protocol, we tested the workflow on the setup shown in Supplementary figure 2(a). The setup consisted of a bottle of room-temperature deionized water and a bottle of heated deionized water (heated in the microwave for 15 seconds before the experiment). Temperature probes were inserted from the top into the center of both bottles. We calculated the thermal coefficient α and presented the temperature calibration equation in Supplementary figure 2(b). We compared our calculated α with previous literature values.

### MRFT and PRFS thermometry comparison experiment

To compare the results of MRFT and PRFS thermometry with the gold standard (GS) measurements, we repeated the same setup in Figure 1(a) but with different bovine muscle samples. The experiment protocol included a TSE structural scan, followed by an SPGR sequence and MRF sequence every five minutes for two hours. We produced the temperature maps using the sensitivity calibration equations described above. We selected ROIs of 25 pixels on Δs maps and T_1_ maps around the location of the fluoroptic probe based on the TSE images and calculated the mean of ROIs. The mean Δs and T_1_ were then converted to temperature using the above-mentioned calibration equations.

To compare the MRFT and PRFS thermometry with GS probe measurements, respectively, we averaged the probe measurements during the period of the MRF and SPGR sequence (18 seconds and 16 seconds respectively). We calculated the root mean square error (RMSE) by comparing the mean fluoroptic probe measurements with MRFT and PRFS thermometry predicted temperature, respectively.

To compare the MRFT and PRFS thermometry predicted temperature because the MRF and SPGR sequences were not run simultaneously, we linearly interpolated the MRFT and PRFS thermometry predicted temperature to compare these two methods at the same time point. We calculated the temperature prediction from both methods every 250 seconds between 1000 and 6500 seconds. Correlation plots were graphed and R^2^ values were calculated for probes 3 and 4 (heated bovine muscle center and periphery). The R^2^ values were not reported for probes 1 and 2 (cold muscle center and room-temperature water) due to the small temperature change ranges.

### DBS lead heating experiment

The setup for the DBS lead experiment is detailed in Figure 1(b). The MR protocol for the lead experiment consisted of the TSE structural scan, an MRF sequence for baseline T_1_ measurements, a TSE sequence with different parameters for heating, and an MRF sequence every minute until the temperature returned close to the baseline temperature. The heating sequence had the following parameters: TR=500 ms, TE=42 ms, flip angle=111°, FOV of 22.4 cm × 22.4 cm, slice thickness=2 mm, matrix size=224×224, number of slices=100, acquisition time=14:33 (min:sec). We repeated the heating cycle three times in one single DBS lead experiment.

We selected regions of interest (ROIs) of 9 pixels on T_1_ maps around the location of the fluoroptic probe based on the TSE images. We calculated the mean of the ROIs and used the calibration equation for probe 1 in the temperature sensitivity calibration experiments to convert T_1_ to temperature.

### B_0_ mapping experiment with DBS lead

Because the orientation of the vendor-supplied gold standard B_0_ maps acquisition is limited to the axial plane, we conducted a separate experiment to investigate the effects of B_0_ inhomogeneity in the lead experiment. The lead was inserted into the cold bovine muscle, shown in Figure 1(c). A vendor sequence (fast GRE-based B_0_ mapping) was run to acquire the GS B_0_ maps. The parameters of the B_0_ mapping sequence were TR=250 ms, FOV=250 mm, slice thickness=5 mm, and FA=30 degrees. The total acquisition time was 2:12 (min:sec). We then ran a MRF sequence to compare B_0_ maps. We selected ROIs of 25 pixels surrounding the tip of the DBS lead and calculated the means.

### ROI analysis

For the temperature sensitivity calibration experiment, four regions of interest (ROI), each with 25 voxels, were selected based on the position of the probes, shown in the TSE structural images. The mean T_1_ of each ROI was calculated. The corresponding probes’ temperatures were averaged over the duration of the MRF acquisition (18 seconds). For the DBS lead experiment, we selected ROIs of 9 pixels that were 1 mm, 3 mm, and 6 mm from the tip of the lead electrode to investigate the spatial temperature gradients near the DBS lead.

### Statistics

We required a regression method robust to noise and outliers as the MRFT method relies on converting T_1_ to temperature, a method that is robust to noise and outliers is desired. Hence, we explored five different linear regression models: least squares regression, weighted least squares regression incorporating variation of averages, least squares regression with geometric means, least squares regression with medians, and Passing-Bablok regression (Passing and Bablok 1983), for generating the calibration equation from the calibration experiment. We computed the mean squared error (MSE) for the five models between model-predicted data and experimental data.

## Results

### Temperature sensitivity calibration

Figure 2(a) shows that the T_1_ decreases as the heated bovine muscle (bottom) cooled down and T_1_ increases as the cold bovine muscle (top) heated up for the first 110 minutes of the experiment calibration experiment. The T_1_ of the room temperature water bottle served as a reference and was consistently at ∼2 seconds during the entire experiment. The cold muscle has longer T_1_ than the heated muscle because the heated muscle loses water during the heating process which we drained (see the color difference between the hot and cold muscle in Figure 2(a)). Figure 2(b) presents the best fit line for the center of cold muscle and the center and periphery of heated muscle and their respective R^2^ statistics using simple linear regression. The x-axis is T_1_ and the y-axis is temperature. All R^2^ values are larger than 0.988, indicating a strong linear relationship between temperature and T_1_. The slopes for probe 3 and probe 4 are very similar (199.1 and 195.4 degrees/second) because they are from the same heated bovine muscle. This linear relationship between temperature and T_1_ agrees with previous literature (Debatin and Adam 2012, Nelson and Tung 1987).

**Figure 2:**
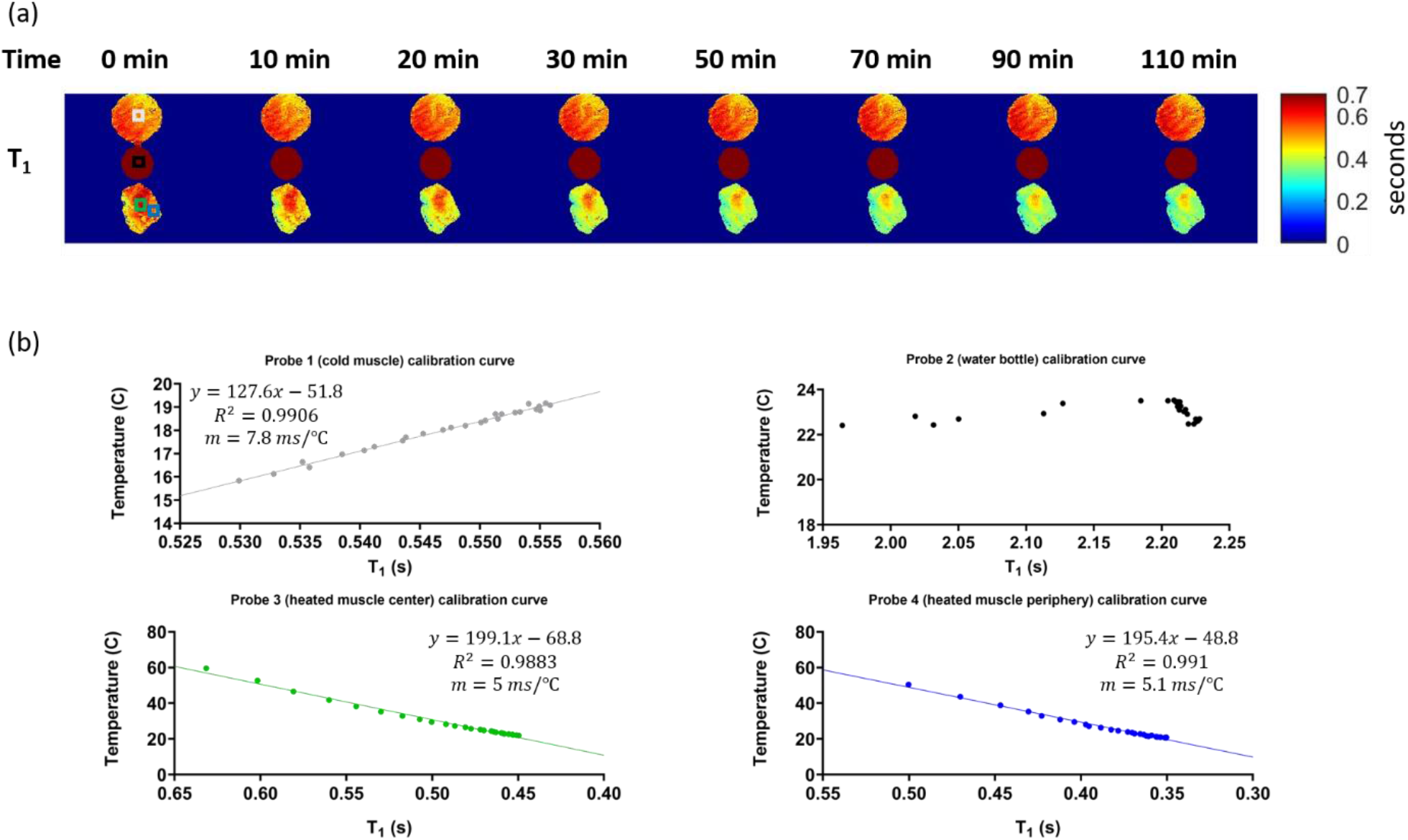
Figure 2 shows (a) a series of T_1_ maps acquired using MRF at the first 110 minutes during the calibration experiment. The time represents the total time from the beginning of the experiment. Regions of interest (ROI) are drawn based on the locations of the probes to calculate the T_1_ estimations around the probes. Each ROI contains 25 voxels. (b) The mean of T_1_ estimations calculated using ROIs over the measured fluoroptic probe temperatures for all probes and the resulting fit using least squared fitting. The best equation for best-fitted line and their R^2^ values are presented in the figure. No fitting was performed for the reference water.

Figure 3(a) shows that Δs increases as the heated bovine muscle (bottom) cooled down and Δs decreases as the cold bovine muscle (top) heated up for the first 110 minutes of the calibration experiment. The room temperature water bottle served as a reference, but the inverse linear relationship between Δs and temperature can also be observed. Figure 3(b) presents the best fit line for the center of cold muscle and center and periphery of heated muscle and their respective R^2^ statistics using simple linear regression. The x-axis is Δs and the y-axis is temperature. All R^2^ values are larger than 0.958, indicating a strong linear relationship between temperature and Δs. The thermal coefficients for probes 3 and 4 are very similar, while the thermal coefficient for probe 1 is higher.

**Figure 3:**
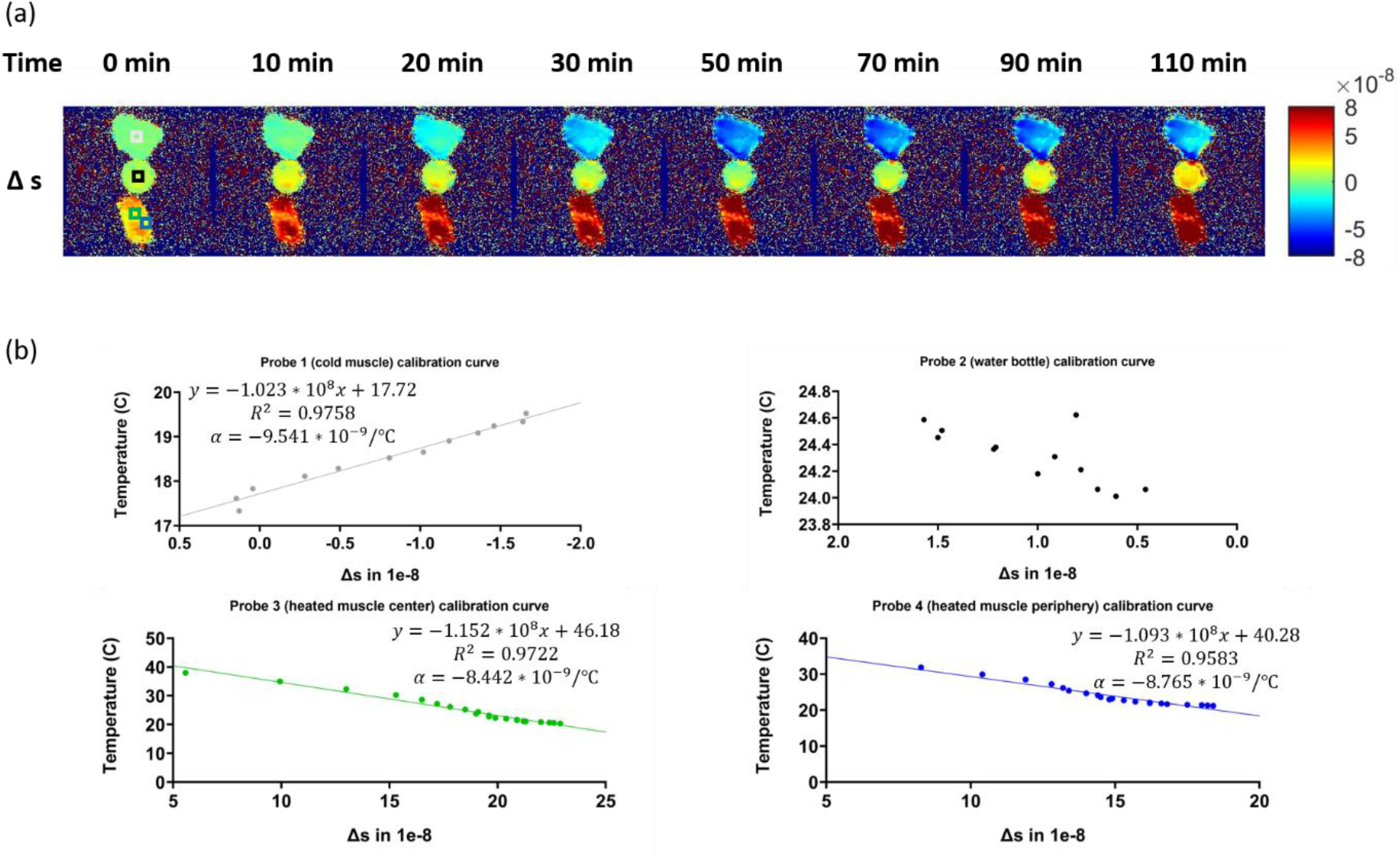
Figure 3 shows (a) a series of Δs maps acquired using MRF at the first 110 minutes during the calibration experiment. The time represents the total time from the beginning of the experiment. Regions of interest (ROI) are drawn based on the locations of the probes to calculate the Δs estimations around the probes. Each ROI contains 25 voxels. (b) The mean of Δs estimations were calculated using ROIs over the measured fluoroptic probe temperatures for all probes and the resulting fit using least squared fitting (the first 12 points were used for curve fitting of probe 1). The best equation for the best-fitted line and their R^2^ values are presented in the figure. No fitting was performed for the reference water.

The five linear regression models and their respective mean squared error (MSE) are shown in Supplementary table 1. For the calibration experiment, we selected the least squares regression because it produces the smallest MSE compared to other models for the cold muscle, which was the same muscle used for the DBS lead experiment.

### MRFT and PRFS thermometry comparison experiment

Figure 4 shows the results of the MRFT and PRFS thermometry comparison experiment. Figure 4(a) shows the comparison of GS probe measurements, PRFS thermometry predicted temperature, and MRFT predicted temperature. Because we ran SPGR sequence before the MRF sequence, MRFT predicted temperature lags PRFS thermometry predicted temperature in time. RMSE of MRFT and probe measurements and RMSE of PRFS thermometry and probe measurements were calculated and presented. Figure 4(b) presents the correlation plots of probes 3 and 4. All R^2^ values are larger than 0.92, indicating a strong linear relationship between MRFT and PRFS thermometry.

**Figure 4:**
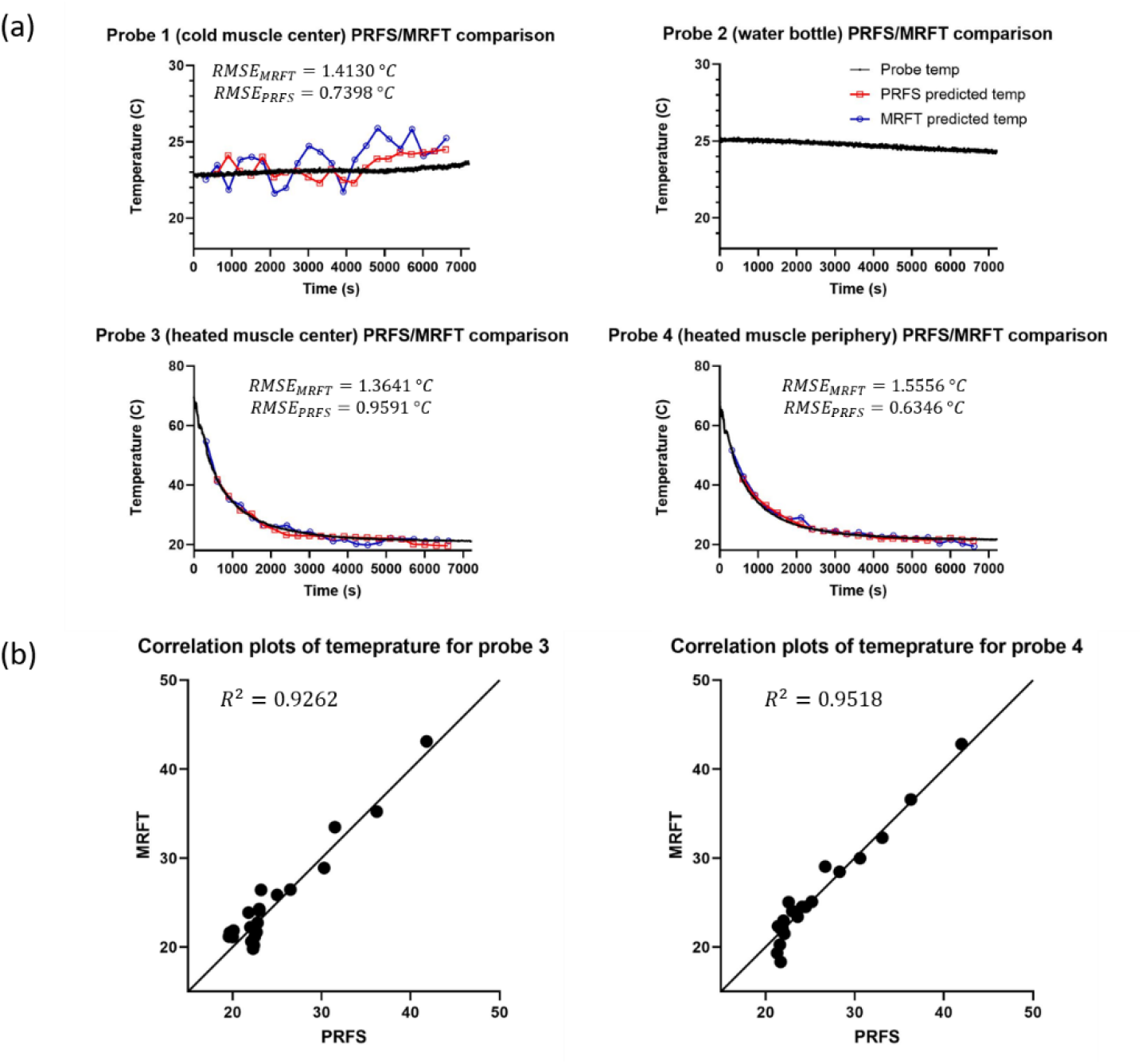
Figure 4 shows (a) the comparison of fluoroptic probe temperature, PRFS predicted temperature, and MRFT predicted temperature for the T_1_-based MRF and PRFS thermometry comparison experiments. The temperature prediction of MRFT and PRFS are shown at the time the sequence started. MRF sequence was run right after the SPGR sequence. The first SPGR sequence served as a baseline and was not shown. Root mean square error was calculated between PRFS and fluoroptic probe temperatures and MRFT and fluoroptic probe temperatures during their sequence time respectively. (b) The correlation plots between PRFS thermometry and MRFT. The temperature predictions of both methods were interpolated to compare their results at the same exact time.

### DBS lead experiment

Figure 5(a) shows the T_1_ map of the first heating cycle in the lead experiment. The T_1_ increases as temperature increases due to TSE-induced heating surrounding the lead (bottom bovine muscle). An ROI was selected in the TSE structure image Figure 5(c) to investigate the temperature estimated by MRFT and the temperature measured by a fluoroptic probe. Figure 5(d) shows that MRFT estimated temperature has a similar trend as the probe temperature but underestimates the values. The blue arrow indicates the start of one heating cycle and the red arrow indicates the end of one heating cycle. The smaller range of MRF predicted temperature compared to the probe temperatures may be because the ROI was 1 mm away from the tip of the lead, which may cause spatial averaging of T_1_ values.

**Figure 5:**
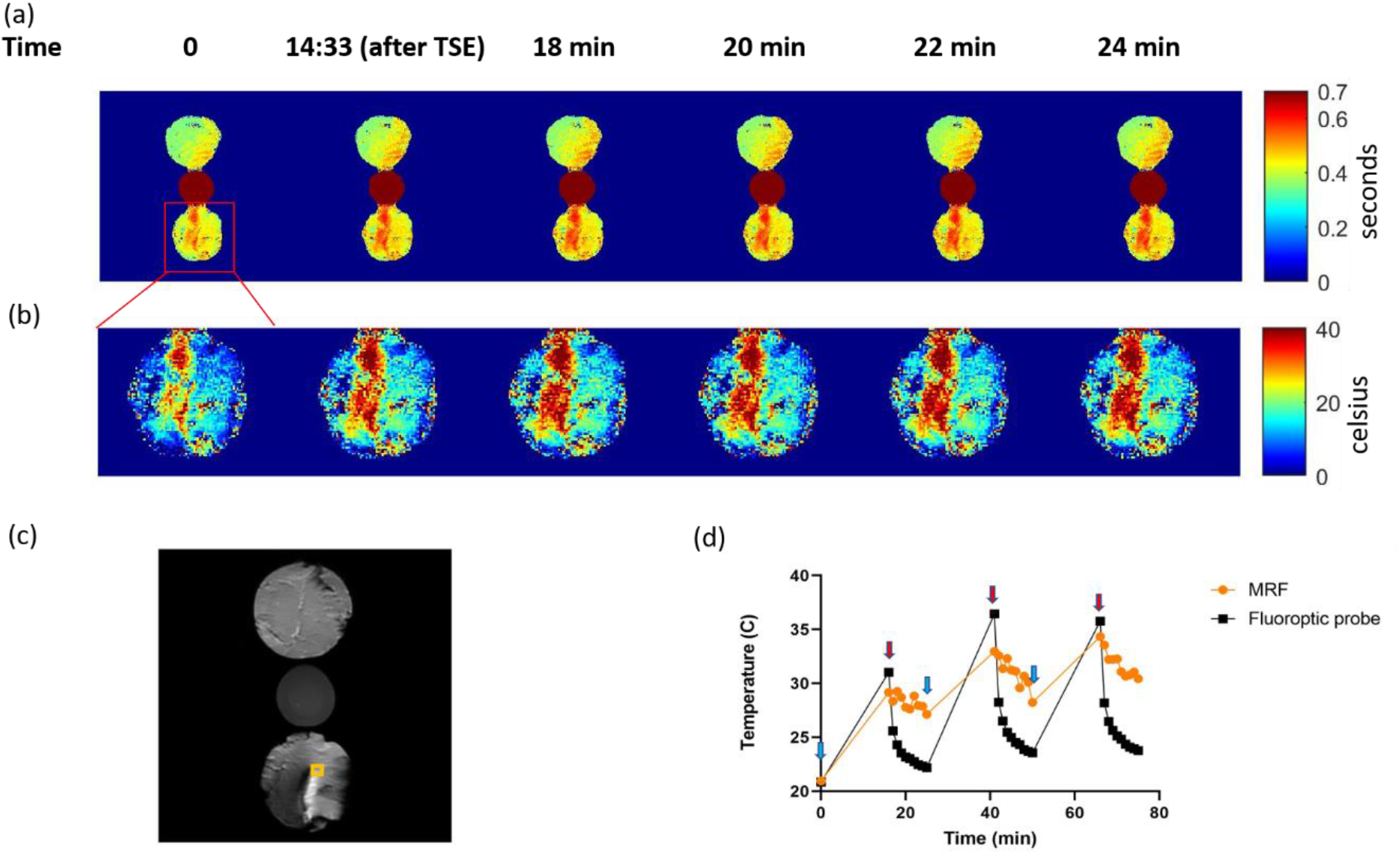
Figure 5 shows (a) a series of T_1_ maps acquired using MRF at the first 24 minutes during the lead experiment. The time represents the total time from the beginning of the experiment. (b) The magnified temperature maps of the bovine muscle with the lead. We used the calibration equation from the temperature sensitivity calibration experiment to convert T_1_ maps to temperature maps. (c) A structure image acquired using TSE at the end of the experiment. We selected the ROI at the tip of the lead corresponding to the position of the fluoroptic probe. Each ROI contains 9 pixels. (d) The comparison of maximum temperature in the ROI estimated using MRF and temperature measured with fluoroptic probe. The baseline for each cycle are marked with blue arrow. The measurements right after the TSE heating are marked with red arrow.

### ROI analysis - Distance from the lead

Figure 6 shows the position of the three different ROIs that are 1 mm, 3 mm, and 6 mm from the tip of the lead and their maximum values for each time point respectively. As expected, the 1 mm ROI has the largest temperature change (∼ 10 °C) and the 6 mm ROI has the smallest temperature change (∼ 4 °C). This is similar to the findings in (Shrivastava *et al* 2012). The starting temperature for the ROIs was different, which may be because the temperature of the muscle was not homogeneous when the experiment started.

**Figure 6:**
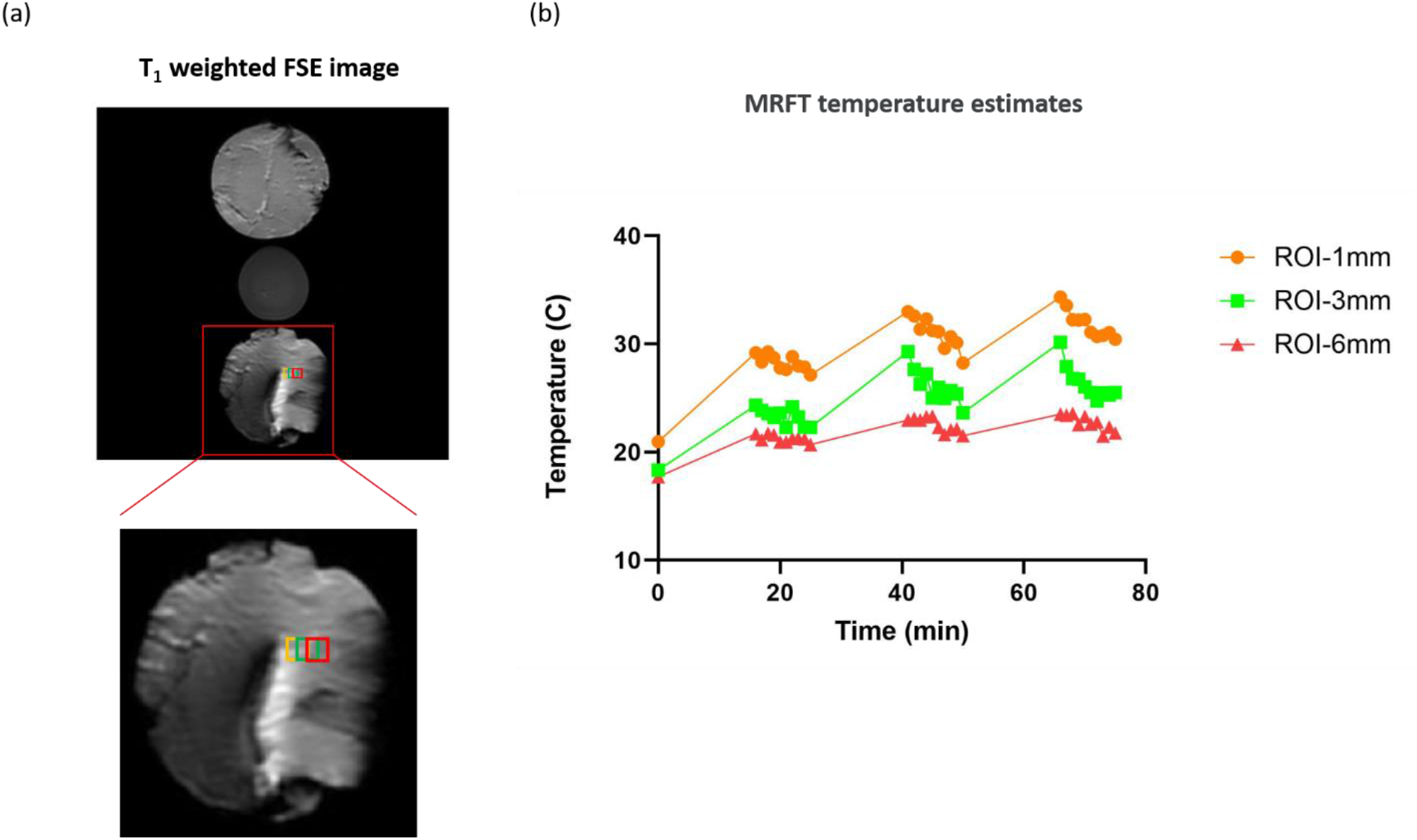
Figure 6 presents three ROIs with 1 mm, 3 mm, and 6 mm distance to the lead. Each ROI contains 9 pixels (3 pixels by 3 pixels square). (a) shows the position of these three ROIs and the figure below presents the zoom-in figures of the region. (b) shows the maximum temperature of the ROIs estimated using MRF.

### B_0_ experiment with DBS lead

Figure 7(a) shows the overlay of TSE structural scan and B_0_ maps generated using vendor-supplied B_0_ mapping sequence and MRF sequence. Both maps show larger B_0_ inhomogeneities around 200 Hz around the tip of the lead where the electrodes were positioned. Figure 7(b) shows that MRF overestimated the B_0_ map by ∼40 Hz in the ROIs we selected. This may be attributed to the fact that vendor-supplied B_0_ mapping sequence utilized Fourier Transform for the phase reconstruction and MRF exploits pattern recognition to generate B_0_ maps. These differences require further investigation.

**Figure 7:**
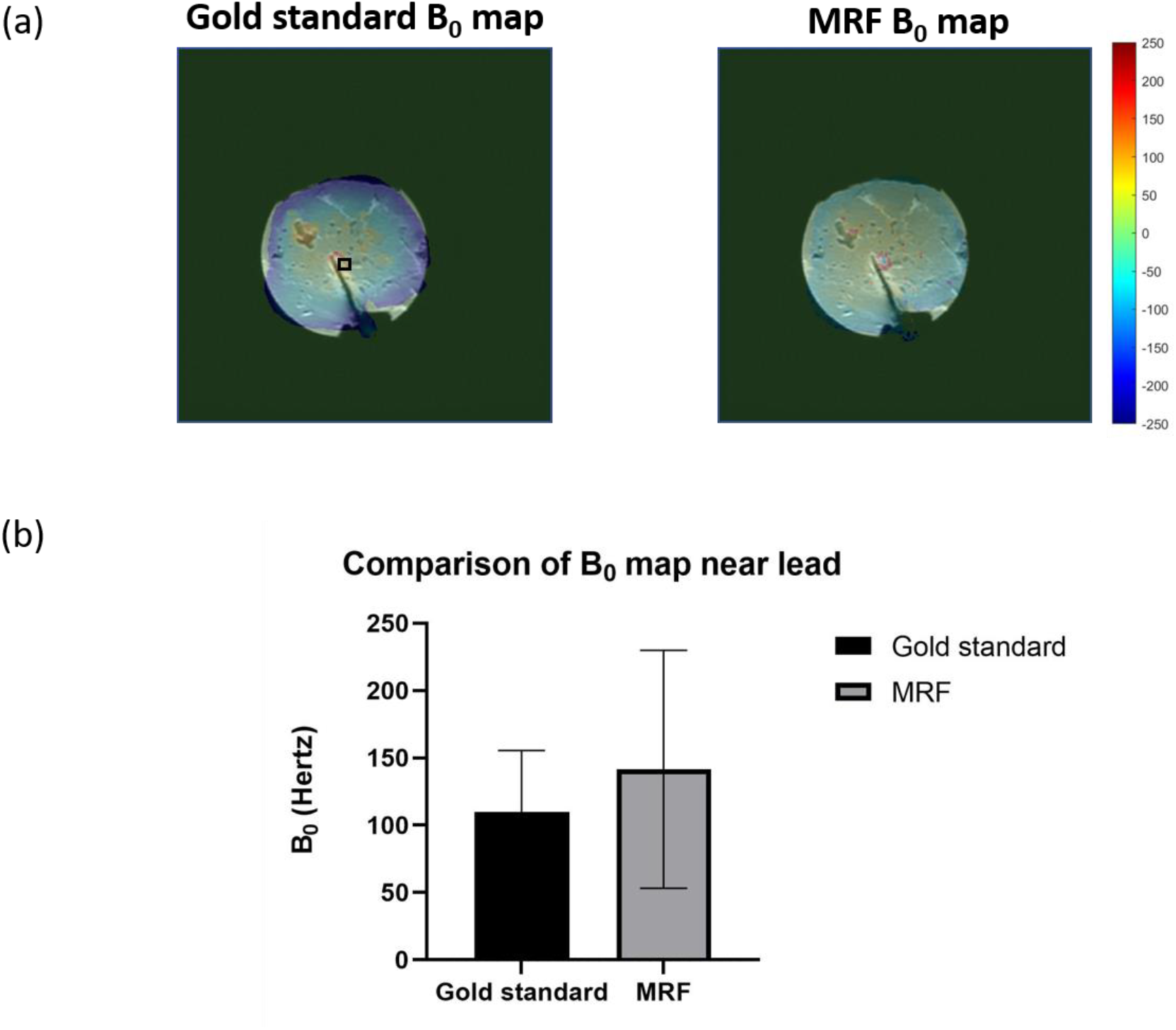
Figure 7 shows (a) the overlay of TSE structure image and B_0_ maps generated using gold standard sequence and MRF. We selected a rectangle ROI of 25 pixels near the lead to compare the values. (b) Comparison of ROI values between gold standard and MRF generated B_0_ maps.

## Discussion

In this study, we present the first work of T_1_-based MR thermometry using MR fingerprinting (dubbed MRFT). We compared MRFT with PRFS thermometry on *ex vivo* bovine muscle and demonstrated its feasibility on *ex vivo* bovine muscles with a DBS lead.

MRFT has several advantages over PRFS thermometry. PRFS thermometry is sensitive to inter and intra-scan motion (Winter *et al* 2016, Odéen and Parker n.d.). Previous studies have shown that MRF preserves structures in parametric maps despite subject motion (Yu *et al* 2018, Ma *et al* 2013). MRFT is also a reference-less method. Once the calibration equation is acquired, no baseline measurement is required to generate absolute temperature maps. MRF can generate multi-parametric maps simultaneously. This provides the potential to perform T_1_-based MRFT, T_2_-based MRFT, PD-based MRFT, and ADC-based MRFT and combine them to provide more robust temperature estimates by simply averaging the outcomes from the different tissue parameters.

Our current implementation of MRFT also has some limitations. The microwave heating we used was uneven and introduced errors in temperature estimation. We also did not test MRFT in ASTM standard gel phantom. MRFT takes ∼30 seconds/slice for temperature estimation. This needs to be sped up to allow real-time temperature estimation.

The position and configuration of the leads have a substantial effect on the electric field and SAR distribution (Shrivastava *et al* 2012, Mattei *et al* 2008, Golestanirad *et al* 2019). These will affect the amount of energy absorbed by tissues surrounding the DBS lead, thus affecting the observed heating in those regions. This is clinically significant, as leads are usually looped several times in a random pattern in conventional DBS surgery (Golestanirad *et al* 2019). For this study, we focused on investigating MRF thermometry and used the same DBS position and configuration for all experiments. We will compare the temperature maps of different DBS positions and configurations in future *in vivo* experiments.

The thermal coefficient α in the PRFS temperature calibration experiment is usually assumed to be around −1 * 10^−8^/°C, but values between −0.7 * 10^−8^/°C and −1.1 * 10^−8^/°C were reported for different tissue types (Odéen and Parker n.d., McDannold 2005). Although PRFS method is regarded as tissue independent to a large degree, because the phase changes are usually very small, on the order of 10^−8^/°C, even tiny discrepancies will lead to large errors in the final temperature maps. In this study, we reported α between −0.84 * 10^−8^/°C and −0.95 * 10^−8^/°C for bovine muscles, as shown in Figure 3, and −10.25 * 10^−8^/°C for DI water, as shown in Supplementary figure 2. These values agree with literature values. The differences in α can be attributed to the inhomogeneous heating during microwaving in our experiment and the temperature-dependent changes in tissue electrical properties.

Temperature dependence between T_1_ relaxation times and the temperature was found to be on the order of 1%/°C (Lewa and Majewska 1980), with values of 1.4%/°C in bovine muscle (Cline *et al* 1994, Rieke 2012). In this study, we reported the temperature dependence of 1.4%/°C for cold bovine muscle center, 1.1%/°C for heated bovine muscle center, and 1.4%/°C for heated bovine muscle periphery at room temperature 20 °C. These values agree with previous literature values. The muscle differences of temperature dependence between heated bovine muscle center and heated bovine muscle periphery may be attributed to homogeneous heating of the muscle samples. A more homogeneous heating using a water bath will likely improve the results.

PRFS thermometry consistently provides better temperature predictions compared to MRFT for all probes, shown in Figure 4(a). T_1_ is sensitive to temperature. At room temperature 20 °C, 1% of change will change the temperature prediction by ∼1 °C, the MRF sequence has to be repeatable for MRFT to predict temperature accurately. Previous MRF repeatability studies have shown that MRF achieved a coefficient of variation (defined as the ratio of the standard deviation to the mean T_1_) below 5% for a large range of T_1_s (Jiang *et al* 2017, Shridhar Konar *et al* 2021, Körzdörfer *et al* 2019, Buonincontri *et al* 2019, Qian *et al* 2022). In case of DBS lead heating, we expect changes of multiple degrees locally (∼4-6 °C, up to ∼10 °C in this study) and hence find MRFT potentially useful.

The current *ex vivo* bovine muscle experiment design does not consider the effects of perfusion. It has been shown in previous studies that fractional regional blood volume of *in vivo* muscle increases with increased temperature (Morvan *et al* 1993). The increased fractional regional blood volume will in turn increase perfusion, and affect the local magnetic field. This will affect PRFS temperature measurements as well as MRF based T_1_ temperature measurements. However, the perfusion induced error is relatively small, as reported in (Gellermann *et al* 2005).

## Conclusion

In this work, we demonstrated the first work of T_1_-based MR thermometry using MR fingerprinting (dubbed MRFT). We observed strong linear relationships between T_1_ and temperature and Δs and temperature for MRFT and PRFS thermometry in *ex vivo* bovine muscle (R^2^>0.988 and R^2^>0.958 respectively). MRFT and PRFS thermometry both accurately estimated temperature in *ex vivo* bovine muscle (RMSE<1.55 °C and RMSE<0.95 °C respectively). MRFT estimated temperature has a similar trend as the probe temperature near DBS lead. B_0_ maps generated using MRFT showed inhomogeneities near tip of the lead, same as gold standard B_0_ maps. This work may serve as a thermometry scout that enable heat monitoring near DBS lead in patients with DBS implant and introduce cooling periods to keep heating below a safe threshold and ensure safety.

