## Supplementary figures and images for "Magnetic Resonance Fingerprinting based Thermometry (MRFT): application to ex vivo imaging near DBS leads"

### Supplementary figure 1

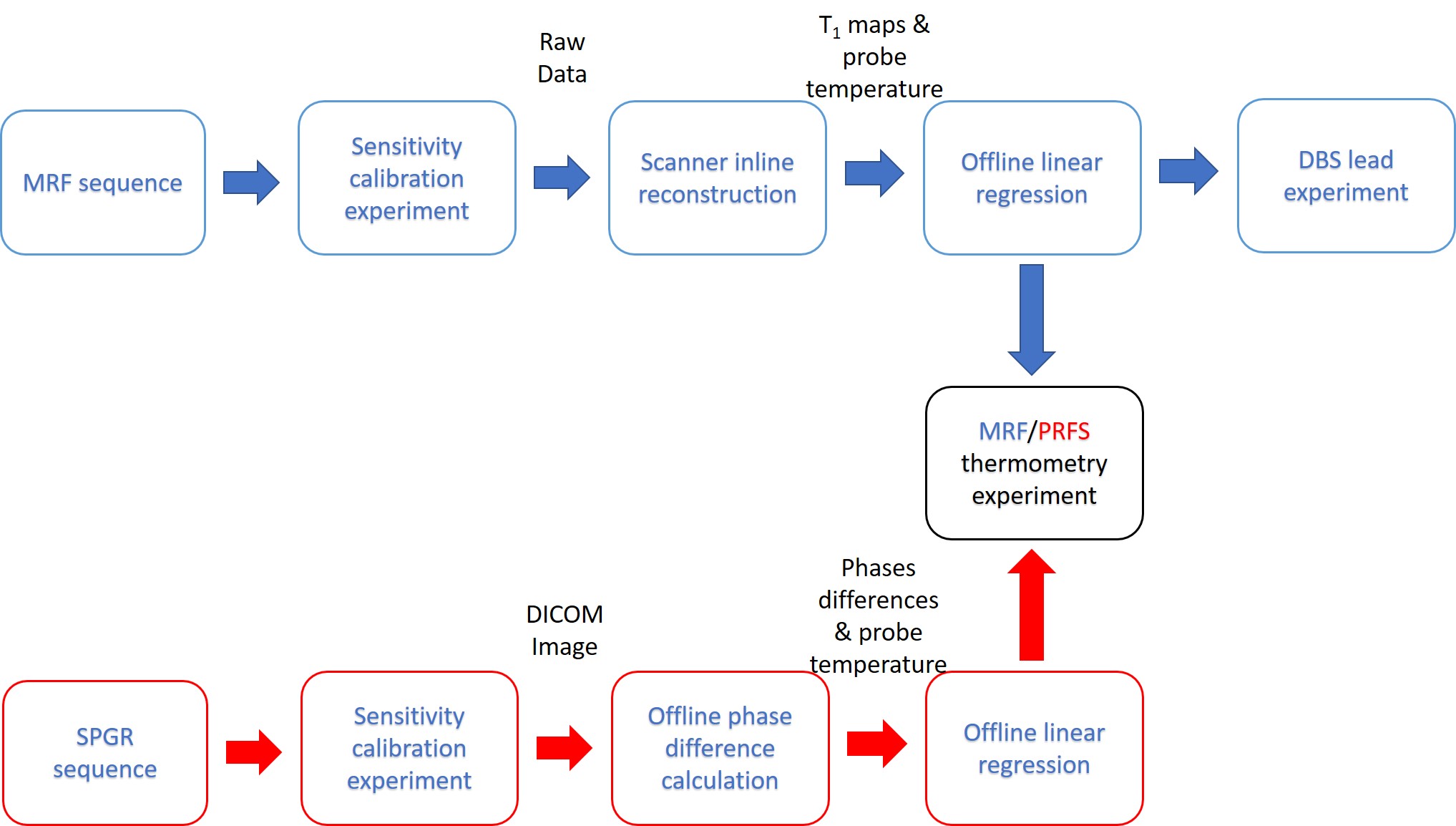

### Supplementary figure 2

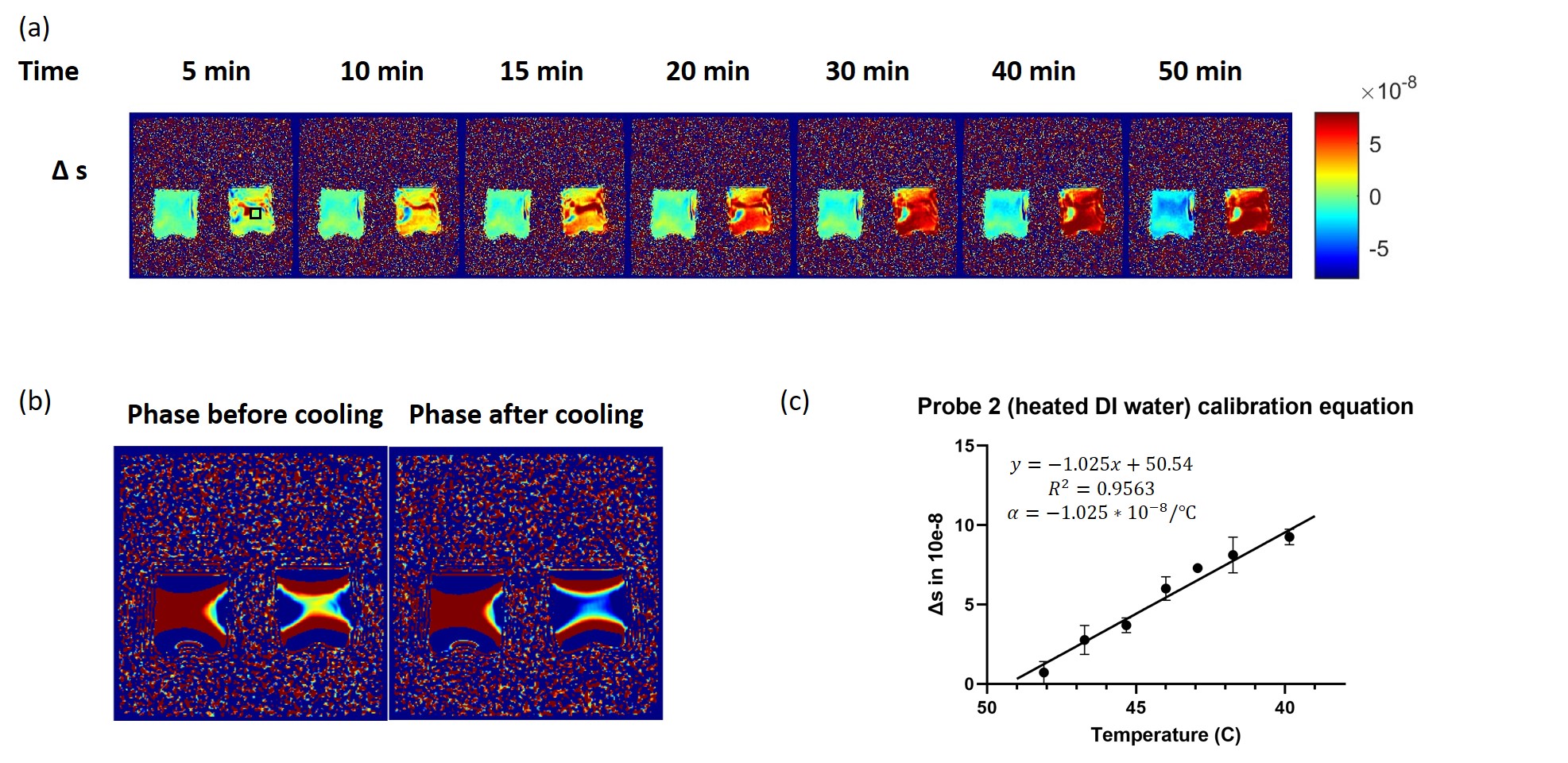

### Supplementary table 1

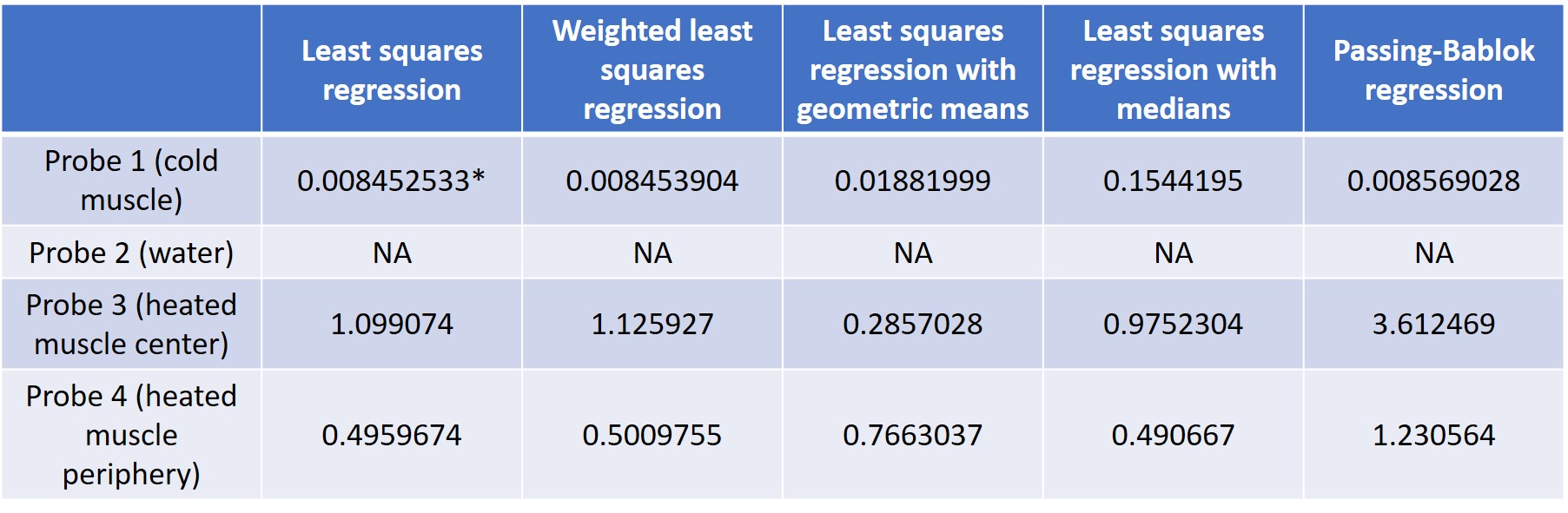
